# Screening and identification of gene expression in large cohorts of clinical lung cancer samples unveils the major involvement of EZH2 and SOX2

**DOI:** 10.1101/2024.09.17.613500

**Authors:** Niharika, Ankan Roy, Ratan Sadhukhan, Samir Kumar Patra

**Author notes:** Correspondence to: **Prof. Samir Kumar Patra,**.

## Abstract

Lung adenocarcinoma (LUAD), the primary subtype of Non-Small Cell Lung Cancer (NSCLC), accounts for 80% to 85% of cases. Due to suboptimal screening method, LUAD is often detected in late stage, leading to aggressive progression and poor outcomes. Therefore, early disease prognosis for the LUAD is high priority. In order to identify early detection biomarkers, we conducted a meta-analysis of mRNA expression TCGA and GTEx datasets from LUAD patients. A total of 795 differentially expressed genes (DEGs) were identified by exploring the Network-Analyst tool and utilizing combined effect size methods. DEGs refer to genes whose expression levels are significantly different (either higher or lower) compared to their normal baseline expression levels. KEGG pathway enrichment analysis highlighted the TNF signaling pathway as being prominently associated with these DEGs. Subsequently, using the MCODE and CytoHubba plugins in Cytoscape software, we filtered out the top 10 genes. Among these, SOX2 was the only gene exhibiting higher expression, while the others were downregulated. Consequently, our subsequent research focused on SOX2. Further transcription factor-gene network analysis revealed that enhancer of zeste homolog 2 (EZH2) is a significant partner of SOX2, potentially playing a crucial role in euchromatin-heterochromatin dynamics. Structure of SOX2 protein suggest that it is a non-druggable transcription factor, literature survey suggests the same; hence, we drove our focus to investigate on potential drug(s) targeting EZH2. Molecular docking analyses predicted most probable inhibitors of EZH2. We employed several predictive analysis tools and identified GSK343, as a promising inhibitor of EZH2.

## Introduction

The epigenome profiles of cancers, particularly lung cancers, are still not well-established. Ongoing research reveals the mechanisms of action of epigenetic modifiers and their interactions with transcription factors and pluripotency inducers. Unfortunately, lung cancer is the second most prevalent cancer in both men and women, and it is the leading cause of cancer-related deaths globally [1]. The International Agency for Research on Cancer (IARC) predicted that lung cancer would cause 1.8 million (18.7%) cancer deaths in 2022 worldwide, with non-small-cell lung cancer (NSCLC) comprising 80% of cases, and despite advancements in treatment, NSCLC survival rates remain dismally low [2–4].

Current research trend highlights the molecular distinctions among histological types of lung cancer, including differences in mutational patterns, DNA copy numbers, aberrant DNA methylation and gene expression signatures. The growing popularity of molecular genetic classification aligns with the need for more targeted approaches [5–8]. For molecular profiling of cancers, next-generation sequencing (NGS) is essential, especially, in combination with RNA-seq analysis, it could rectify the drawbacks of microarray analysis and enable precise identification of low-abundance transcripts. Integrating microarray technology and bioinformatics, a high-throughput approach for unveiling genetic alterations has found widespread application in investigating the mechanisms behind tumor development and cancer progression. This combined methodology has proven invaluable in identifying novel diagnostic and prognostic biomarkers for various cancers [9,10]. A publically available database, NCBI-Gene Expression Omnibus (GEO) provide copious amounts of data for scientific pursuits [11]. Numerous studies have utilized data mining techniques on the GEO database through bioinformatics analyses to pinpoint DEGs in multiple cancers.

More than 1600 transcription factors (TFs) involved in human transcriptional network and are highly regulated. Sry-related high-mobility box 2 (SOX2), a key pluripotency-associated stem cell factor, is crucial for maintaining stem cell properties and determining cell fate, thus regulating developmental processes [12]. SOX2 had been detected to be abnormally expressed in many cancers, including lung, breast, colon, ovary, and prostate carcinomas, with its expression positively correlated with cancer cell stemness and poor patient outcomes, indicating its significant role in cancer stem cell generation and biology [13–16], for a comprehensive review, see Niharika et al., 2024 [17]. Recent genomic studies have revealed that many chromatin regulators play crucial roles in cancer development, and EZH2, a histone methyltransferases dedicated to methylate H3K27 is the subunit of a Polycomb Repressor Complex 2 (PRC2). EZH2 recurrently mutated in several cancers and highly expressed in many others [18–20]. EZH2 forms complexes with transcription factors or directly binds to the promoters of target genes to regulate gene transcription; hence, it has become a prominent target for cancer therapy, leading to the development of an increasing number of potential medicines. Dysregulation of EZH2 has been observed in various caner types including breast cancer [18], Non-Small Cell Lung Cancer [19], gastric cancer [20], and ovarian cancer [21]. However, the SOX2-EZH2 axis has not yet been reported as therapeutic target in any types of cancer; we are projecting that this could be a promising diagnostic approach and an attractive therapeutic target to eliminating CSCs and overcoming chemoresistance.

From the GEO database (http://www.ncbi.nlm.nih.gov/geo/) [22], we have acquired GSE19188 and GSE68465 datasets. The online tool Gene Expression Omnibus 2R (GEO2R) was employed to identify DEGs between LUAD and normal lung tissue clinical samples in the above-mentioned datasets. Commonly up and downregulated genes were identified using Venn diagram software. We next performed functional enrichment analysis, which included Kyoto Encyclopedia of Genes and Genomes (KEGG) pathway analyses and gene ontology (GO), followed by protein-protein interaction (PPI) network construction. We used several databases and tools, including The University of Alabama at Birmingham CANcer (UALCAN, http://ualcan.path.uab.edu/index.html) data analysis web tools, Kaplan-Meier plotter [KM plotter, (http://kmplot.com/analysis)], Gene Expression Profiling Interactive Analysis 2 [GEPIA2, http://gepia2.cancer-pku.cn/], and Human Protein Atlas (HPA) database (https://www.proteinatlas.org/), to validate the selected candidate genes. Gene-TFs interaction analysis and their correlations guided us in narrowing down target sets for therapeutic purposes. Employing bioinformatics-based research tools we aimed to identify potential biomarkers for LUAD.

We identified a new set of genes (hub genes) responsible for LUAD progression through gene expression and pathway analyses, and protein-protein network studies. Notably, a direct correlation between the transcription factor SOX2 and the histone modifier EZH2 was observed, and experimental verification of SOX2 expression ensured that, EZH2 does not impose any repressive burden on SOX2 gene promoter. Additionally, from the available information by literature survey it is apparent that, that EZH2 repressed/silenced many tumor suppressor genes (TSGs) such as ARH1, TIMP2, GPER1, MASPIN, and SLC17A7 in ovarian cancer, breast cancer, non-small cell lung adenocarcinoma (NSCLC), and glioblastoma stem cells, respectively, by by trimethylation of histone 3 lysine 27 (H3K27me3) in the gene promoter [23–27]. We are the first to find out that SOX2 is highly expressed among the 10 DEGs in the context of hub gene analysis in the LUAD. However, there is no report on the interaction of EZH2 with SOX2 as of yet. It is well known that SOX2 expresses in combination and coordination with other specific transcription factors to revert the terminally differentiated cells to a more potent lineage stage; such as OCT4, KLF4, and c-MYC in one group and OCT4, NANOG, and LIN28 in another [28,29]. Based on our results we hypothesize that, when SOX2 interacts with EZH2, it would silence the gene in LUAD or reversely EZH2 fails to represses genes when SOX2 interacts with it.

## 2 Materials and method

### 2.1 Microarray data information

Considering the GEO database, we searched for “Lung Cancer” and specified “Homo sapiens” under “Top Organisms” and “Expression profiling by array” for the “Study type.” The inclusion criteria were: (a) Every dataset includes a control sample group (healthy lung tissues) and a LUAD sample group; (b) each group includes a minimum of two samples. To eliminate exclusion criteria were used (a) duplicate studies, (b) letters, (c) review articles, (d) case reports, (e) incomplete data, (f) non-human studies, and (g) datasets lacking a control sample group. Following the above-mentioned criteria, only two microarray expression profile datasets, GSE68465 [30] and GSE19188 [31] were downloaded.

### 2.2 Data acquisition and processing

To get the DEGs, GEO2R, a web-based tool was used to compare between the control (normal lung samples) and test (LUAD samples) groups. The identification of key regulators/genes with distinct expressions was done using p-value measurements as well as fold change (FC) statistics. A p-value criterion of less than 0.05 was used. In a case-control analysis, FC, a widely employed method in differential expression analysis, quantifies the extent of gene expression changes between two conditions, such as tumor and normal samples. We found common up- and common downregulated genes among the DEGs using the Venn Diagram from the R package. After that, additional downstream analysis was performed on these common genes.

### 2.3 Enrichment analyses of common DEGs

All differentially expressed genes (up and downregulated) underwent a thorough examination through GO and KEGG pathway analyses [32–34]. GO enrichment analysis, a method for interpreting gene sets based on their functional characteristics was carried out in this study using the Enrichr tool. The ToppFun functional annotation tool was utilized to validate the GO analysis and obtain a thorough comprehension of biological data, which included molecular functions (MF), cellular components (CC), and biological processes (BP) [35]. The statistical significance level was established at p-value < 0.05. Moreover, the KEGG Orthology-Based Annotation System (KOBAS 3.0, http://kobas.cbi.pku.edu.cn/) was used to validate the KEGG pathway enrichment analysis results [36]. The false discovery rate (FDR) < 0.01 and the p-value < 0.05 were established as the significance thresholds.

### 2.4 PPI network and significant module analysis

Establishing gene regulatory networks is crucial for subsequent topological analysis, module identification, and identifying highly influential genes (candidate genes/ hub genes) that may serve as potential targets for cancer prognosis. We used the publicly available Search Tool for the Retrieval of Interacting Genes (STRING) database (https://string-db.org/) to reconstruct the gene regulatory network [37]. As a precomputed global resource, STRING makes it easier to explore and analyze the functional relationships between proteins, which are frequently deduced from the genomic affiliations of the encoding genes. The list of discovered gene biomarkers was uploaded to STRING for interaction retrieval, with the default parameter values being used. After that, Cytoscape 3.10.1 was used to visualize and perform topological analysis on the interaction network that was derived from STRING [38].

### 2.5 Network module identification

Identifying functional units, referred to as network modules or biological pathways, presents a crucial challenge in analyzing intricate and extensive biological networks. Due to the typically vast and intricate nature of biological networks, analyzing them as a whole is impractical, making module identification a pivotal step in gaining biological insights. In biological networks, module identification, also known as community discovery or graph clustering, can be carried out using various software tools and packages created specifically for this purpose. In our approach, we utilized the Molecular Complex Detection (MCODE) [39] clustering algorithm along with the CytoHubba plugin [40]. The CytoHubba plugin offered 11 different topological analysis methods, including six centralities based on shortest paths (Closeness, Betweenness, Bottleneck, Radiality, Eccentricity, and Stress) and Edge Percolated Component, Degree, Density of Maximum Neighbourhood Component, Maximal Clique Centrality, and Maximum Neighbourhood Component. Out of all of these techniques, the most recently suggested MCC technique has demonstrated better results in identifying critical proteins in the human Protein-Protein Interaction (PPI) network using Cytoscape software. Using these techniques, we were able to pinpoint highly coupled areas of the network, which frequently corresponded to pathways and protein complexes.

### 2.6 Validation of candidate genes

The Kaplan–Meier univariate approach with the survival package was used in the survival study to investigate the relationship between candidate genes and patient overall survival (OS). The association between gene expression and survival outcomes in LUAD was examined using the KM plotter. This web-based tool offers a rapid assessment of LUAD prognosis-related genes. The ‘survival’ package was employed to compute and display Kaplan–Meier survival curves. Patients included in the study completed the follow-up period and were categorized into two groups based on the median expression value of the hub gene. A log-rank test with p-value < 0.05 was considered statistically significant.

Using information from hundreds of samples from the Genotype-Tissue Expression (GTEx) and Cancer Genome Atlas (TCGA) projects, the GEPIA2 database [41,42] was used to validate the mRNA expression patterns of potential genes between normal and malignant tissues. Parallel to this, a query was made to the HPA database, and information was taken out to compare the expression of potential genes in cancer specimens and healthy specimens.

### 2.7 A functional network of SOX2 target genes and TFs that regulate SOX2 expression

The operation of multiple downstream genes is influenced by the expression of SOX2 genes, which can also be influenced by other upstream genes. We used the TRRUST database [https://www.grnpedia.org/trrust/) [43] to investigate this regulatory network, which includes data on TFs and the target genes. To find the target genes that these TFs regulated, a subset of TFs was included in the database. As molecular drivers, TFs are essential because they affect the behavior of context-dependent genes and participate in the development of several human cancers. To build this regulatory network, we used NetworkAnalyst [44], a powerful web-based application for statistical, visual, network, and meta-analytical analysis of expression data. The TF-gene regulatory network was built using three current databases: ENCODE, JASPAR, and ChEA. These databases were used to find common transcription factors, and cBioportal was used to conduct co-expression research.

### 2.8 Cell culture

For the validation of our study, two cell lines were utilized. HaCaT, immortalized human keratinocytes, served as the control, while the A549 cell line, and NCI-H522 are human lung cancer cell lines was used. A549, and HaCaT cells were grow in DMEM (Gibco, Cat # 12100046) media and NCI-H522 cells were grow in RPMI 1640 (Gibco, Cat # 31800022) supplemented with 10% fetal bovine serum (FBS, Gibco), 100 units/ml of penicillin, and 0.1 mg/ml of streptomycin (Himedia, SKU: A002A) at 37 °C in a humidified atmosphere with 5% CO2. Prior to seeding for subsequent experiments, the cells were harvested through trypsinization (Trypsin-EDTA 0.25%, phenol red, Gibco, Cat # 25200072), and viable cells was determined via trypan blue staining (0.2% v/v, Gibco, Cat # 15250061) using a hemocytometer.

### 2.9 qRT-PCR analysis

Following the manufacturer’s instructions, total cellular RNA was extracted from each cell line using the TRIzol (Life Technologies, 15596018) technique. Isopropanol (Himedia-MB063) and 70 % alcohol (Himedia-MB228) were used to precipitate RNA and clean off the contaminants, respectively. Then, cDNA was made using 1 μg of total RNA using GeneSure First Strand cDNA Synthesis Kit (Pure-gene, Genetix). Next, quantitative real-time PCR (qRT-PCR) was carried out using PowerUp^TM^ SYBR^TM^ Green Master Mix (Applied Biosystems™, Cat # A25742) in the Agilent Technologies MX300SP Real-time PCR equipment. The housekeeping gene β-Actin was used as an endogenous control to normalize the mRNA levels, and the 2^−ΔΔCt^ technique was employed for the computations [45] (Table 2 containing primer list).

### 2.10 SOX2 protein expression validation using western blotting

Once the cells were reached up to 80–85% confluency, the whole cell lysate were prepared using radio-immunoprecipitation assay (RIPA, sc-24948, Santa Cruz Biotechnology). Subsequently, the Bradford method was employed to determine the protein concentration and immunoblotting procedure was carried out using a modified version of our earlier publication [46]. Equal amounts of cell lysate were loaded and resolved on 8-12% SDS-PAGE gel, depending on the size of protein of interest, in the running buffer. Afterward, proteins were transferred onto a nitrocellulose membrane (Axiva Sichem Biotech Cat # 160300RI). The membranes containing proteins were blocked in 3% BSA in PBS with 0.1% Tween 20 (PBST) for 2 hours at room temperature, followed by overnight incubation with primary antibodies against anti-SOX2 (Abcam-ab59776) and anti-β-Actin [Primary-Mouse IgG1 (sc-47778, Santa Cruz Biotechnology) at 4°C. Subsequently, the membrane underwent three washes with PBST before being incubated with the appropriate horseradish peroxidase (HRP)-conjugated secondary antibody (goat anti-mouse IgG-HRP, sc-2005, Santa Cruz Biotechnology) for 2 hours at room temperature. After thorough washing with PBST buffer, the membranes were developed using the SuperSignal™ West Pico PLUS Chemiluminescent Substrate (Thermo Scientific, Cat #34580) using Bio-Rad ChemiDoc™ MP Imaging System to detect HRP on immunoblots. β-actin used as a control for ensuring equal protein loading. ImageJ quantification software (https://imagej.net/ij/) was used to analyze the relative protein expression obtained from immunoblots.

### 2.11 Protein-inhibitors interactions analysis

#### 2.11.1 Protein preparation

The RCSB Protein Data Bank (PDB, www.rcsb.org) provided the high-resolution crystallographic structure of EZH2, and the PDB ID is 5HYN. The AutoDock 4.2.6 (https://autodock.scripps.edu/), MGL Tools 1.5.6 (https://ccsb.scripps.edu/mgltools/) software packages were used to perform molecular docking studies, and the PyMol 3.0 software package was used to analyze the data. Using AutoDock 4.2.6, polar hydrogen atoms were added; water molecules and inorganic charges were eliminated from the protein. The macromolecule’s charge (Gasteiger) was introduced, and AutoDock 4.2.6’s Lamarckian genetic method was used for ligand docking. To find the binding site, the grid size of x, y and z axes were adjusted to 126, 126, and 126 respectively with a spacing of 1 Å along with center orientation of grid include axes x, y, and z were 84.657, 103.4, and 93.923 respectively. Ultimately, from a pool of distinct conformers, the conformers with the best binding energy were chosen for each docking simulation, and the resulting data was subjected to further analysis. Various parameters were set to their default settings in compliance with the AutoDock 4.2.6 software.

#### 2.11.2 Ligand preparation

The SDF format of all the drug’s three-dimensional (3D) structures was obtained from the PubChem database (https://pubchem.ncbi.nlm.nih.gov/) and then converted to PDB file format. After that, coordinate data were stored in an expanded PDB format called PDBQT using AutoDock 4.2.6, which included atomic partial charges of the atoms and their types. Next, torsion angles were computed to identify molecules’ fixability and nonbonded rotations.

#### 2.11.3 Determination of ligand binding site

FTsite, an open-source server, was used to identify every possible binding site on the EZH2 receptor (http://ftsite.bu.edu/cite) [47]. After uploading the 3D structure with PDB ID 5HYN to the server, a task name was given for result identification. This server utilizes fast Fourier transforms to compute the energy of chemical probes (functional groups) on a grid. This involves identifying clusters formed by each probe individually and assessing the overlap of these clusters among probes to pinpoint potential binding sites. FTSite ranks the detected pockets based on the number of amino acids in contact with the probes but does not provide a quantitative assessment of pocket druggability. As predicted by FTSite, the grid box encompassed the main active sites, with all settings set to default. Later, in order to verify our result, we used the CASTp server to analyze surface topography of the receptor protein. [48].

#### 2.11.5 Protein-drug docking study

EZH2-drug interactions were studied with the AutoDock 4.2.6 software application. The corresponding proteins and drugs docked structures were then visualized using the PyMol and BIOVIA Discovery Studio program.

### 2.12 Statistical analyses

The best statistical technique specified in the databases was followed when downloading the study’s data. An unpaired sample t-test was used to analyze the microarray data retrieved from the GEO databases. Three sets of experiments were carried out for the qRT-PCR and western blot to determine the p-value and validate the findings. The findings were presented as mean ± labeled standard deviation, with a p-value < 0.05 signifying statistical significance.

## 3 Results

### 3.1 Identification of DEGs

We have identified 2427 DEGs (from GSE19188); among those, 938 genes were upregulated, and 1489 genes were downregulated. However, from the GSE68465 dataset, 2988 DEGs were identified. Among those, 1597 genes were upregulated, and 1391 genes were downregulated (Figure 1A & B). Following this, 337 commonly upregulated and 458 commonly downregulated DEGs from both datasets were identified in LUAD samples compared with non-cancerous samples using the Venn diagram software (Figure 1C).

**Fig. 1.**
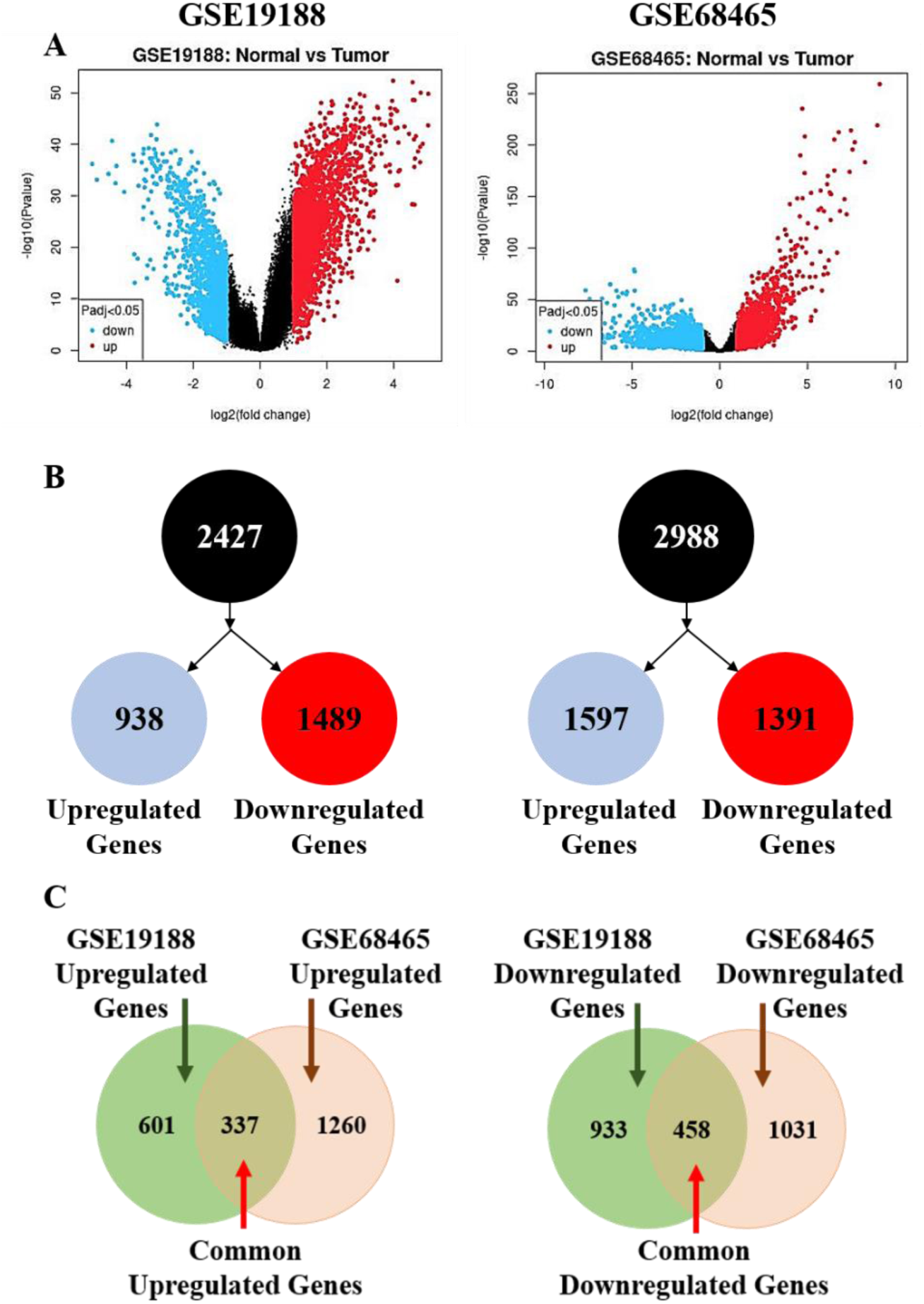
Expression profiles of differentially expressed genes in lung adenocarcinoma. (A) Volcano plot of all genes. Red dots indicate genes that were upregulated blue dots indicate downregulated genes. Each dot represents the expression values of the significant genes. (B) Simplified and counting of differentially expressed genes from two GEO datasets and each datasets having upregulated genes and downregulated genes. (C) The Venn diagram identified total 795 DEGs, including 337 upregulated and 458 downregulated genes are common among these two datasets.

### 3.2 Enrichment analyses

An enrichment analysis using DAVID resulted in 795 DEGs including both types upregulated and downregulated. The GO analysis of common upregulated DEGs considering top biological processes suggested that these genes were involved in Mitotic Sister Chromatid Segregation (GO: 0000070). The cellular components of common upregulated DEGs involved the Spindle (GO: 0005819) and Nucleus (GO: 0005634), while molecular function involved Single-Stranded DNA Helicase Activity (GO: 0017116) (Figure 2A). The GO analysis of common downregulated DEGs in consideration with top biological processes suggested that genes were involved in inflammatory response (GO: 0006954); precisely chosen based on the p-value. The cellular component of common upregulated DEGs involved Collagen-Containing Extracellular Matix (GO: 0062023), while molecular function involved Chemokine Receptor Binding (GO: 0042379) (Figure 2B). Additionally, while we performed KEGG pathway enrichment analysis of all DEGS, the TNF signaling pathway covers maximum DEGs, suggested that TNF signaling gets activated while these DEGs were in combination (Figure 2C).

**Fig. 2.**
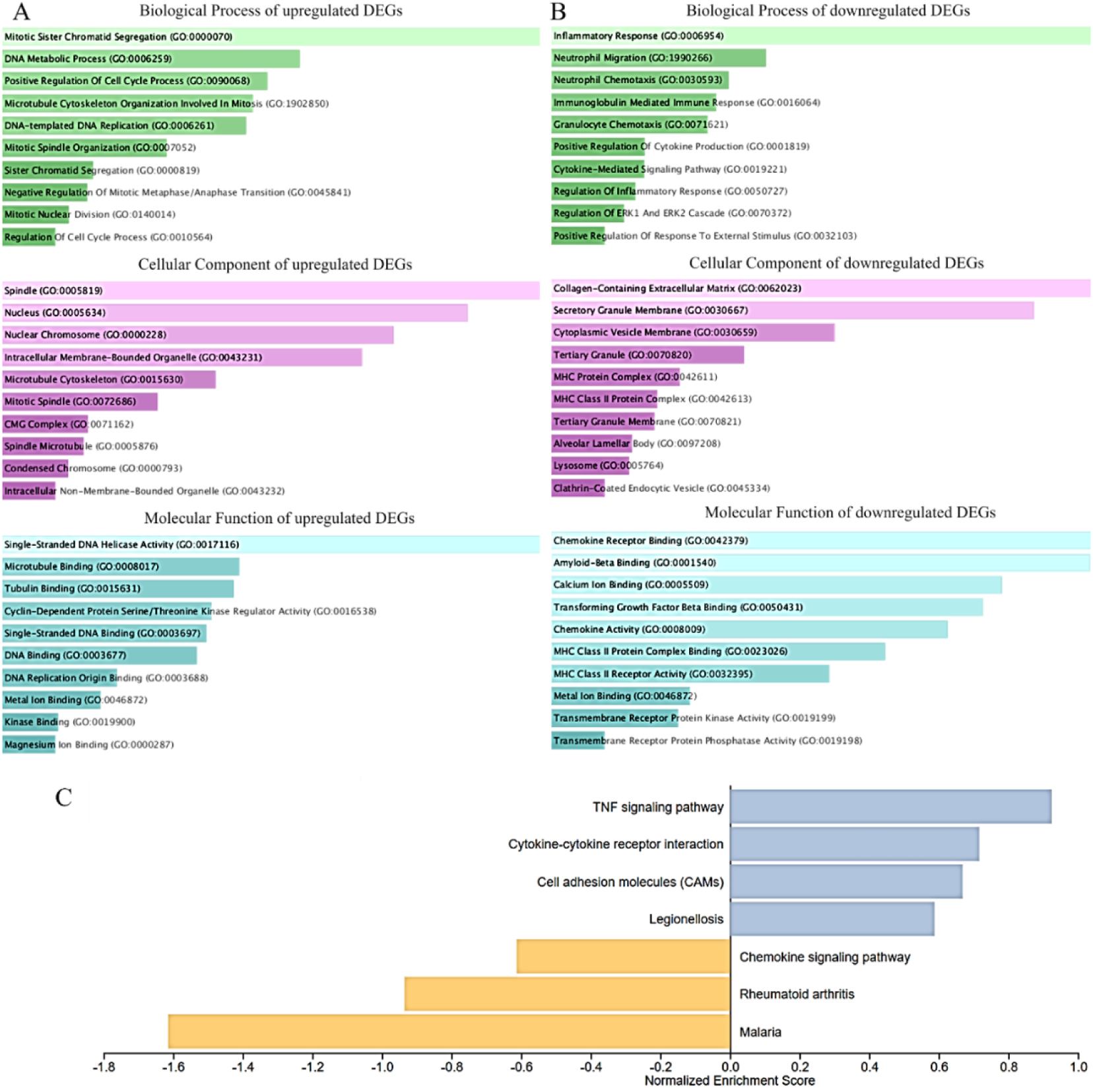
Gene Ontology (GO) term enrichment analysis. Significantly enriched gene set were selected based on a FDR < 0.05. (A) Bar charts showing the top 10 gene ontology terms for common upregulated genes with biological process, cellular components, and molecular function. (B) Bar charts showing the top 10 GO terms for common downregulated genes with biological process, cellular components, and molecular function. (C) Enriched pathways identified using KEGG pathway analysis of DEGs.

### 3.3 Network construction

We build the regulatory network of 795 newly discovered DEGs using the STRING database (Figure 3A). There were 773 nodes and 13256 edges, an average node degree of 34.3, and an average local clustering coefficient of 0.472. PPI enrichment p-value was 1.0e-16.

**Fig. 3.**
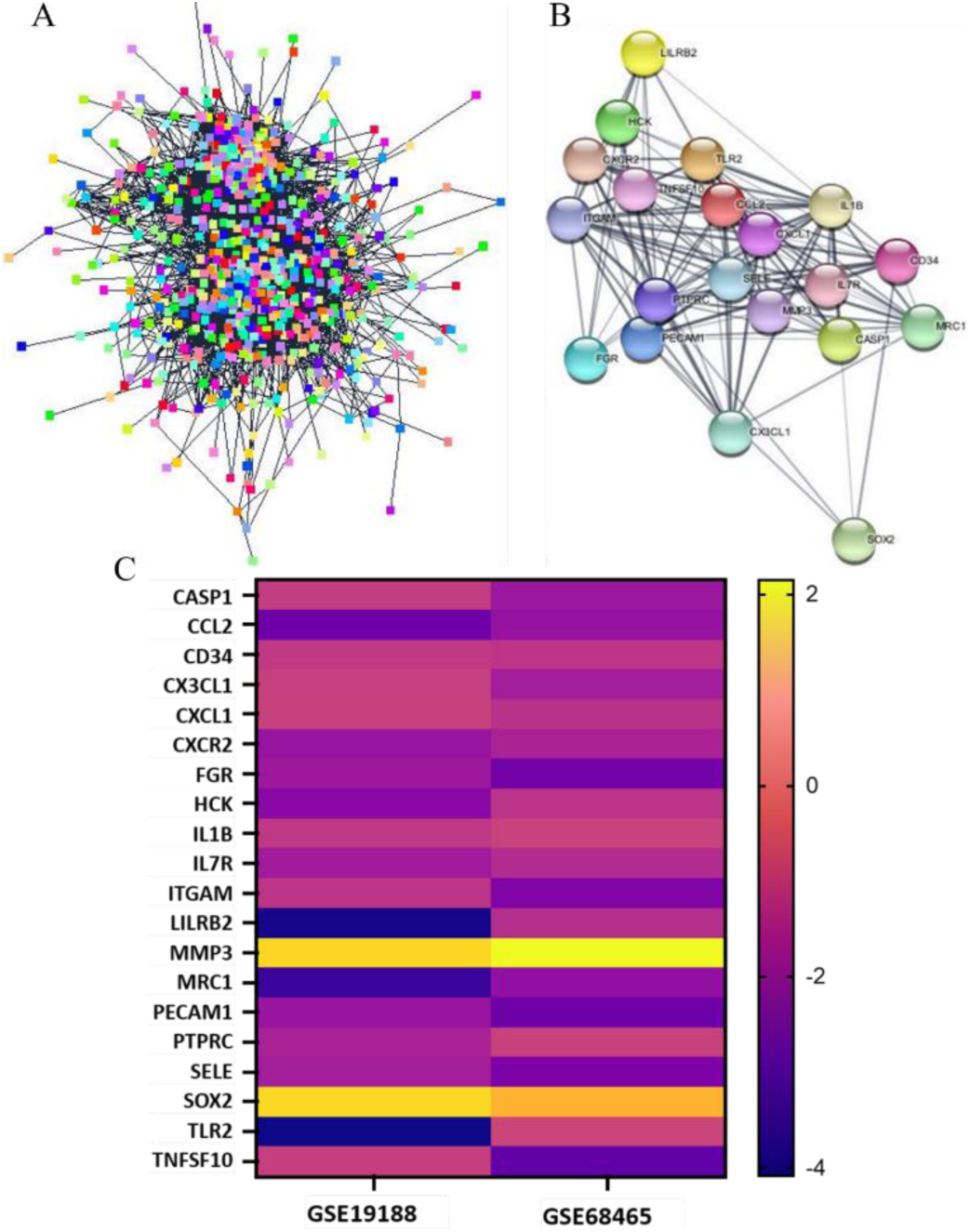
An interactive network of candidate hub genes. (A) Hub genes are interactive with each other within an interactive module. (B) Top 20 hub genes predicted from the DEGs. These hub genes were representative genes involved in occurrence and progression of LUAD. (C) Heatmap of top 20 genes. Blue corresponds to lower gene expression and yellow to higher gene expression.

### 3.4 Module identification in the LUAD network

The MCODE network clustering method was run using its default parameters (degree cutoff = 2, node score cutoff = 0.2, haircut = checked, fluff = unchecked, k-score = 2, maximum depth = 100) and reconstructed the network. A total of fourteen clusters were detected. We selected the top four modules for additional examination. Finally, the top 20 genes were selected by using the CytoHubba algorithm. The top 20 genes include CASP1, CCL2, CD34, CX3CL1, CXCL1, CXCR2, FGR, HCK, IL1B, IL7R, ITGAM, LILRB2, MMP3, MRC1, PECAM1, PTPRC, SELE, SOX2, TLR2, and TNFSF10 (Figure 3B & C).

### 3.5 mRNA and protein expression validation of candidate genes using online databases

As previously stated, survival analysis was performed for the list of genes found exploiting module discovery methods. The Kaplan–Meier univariate survival analysis suggested that out of 20 genes, 12 genes were significant based on p-value <0.05 (Figure 4A). Then, we analyzed the candidate genes’ mRNA expression in tumor and normal samples, and found that 10 genes - CASP1, CX3CL1, FGR, IL7R, ITGAM, MRC1, PECAM1, PTPRC, SOX2, and TLR2 are significant (p-value<0.05, Figure 4B, (description of top 10 genes were mentioned in Table 1). Next, all 10 candidate gene’s protein expressions were analyzed using the HPA database, and it was found that out of 10, 5 genes (CASP1, CX3CL1, FGR, ITGAM, and SOX2) precisely followed their mRNA expression trend (Figure 5).

**Fig. 4.**
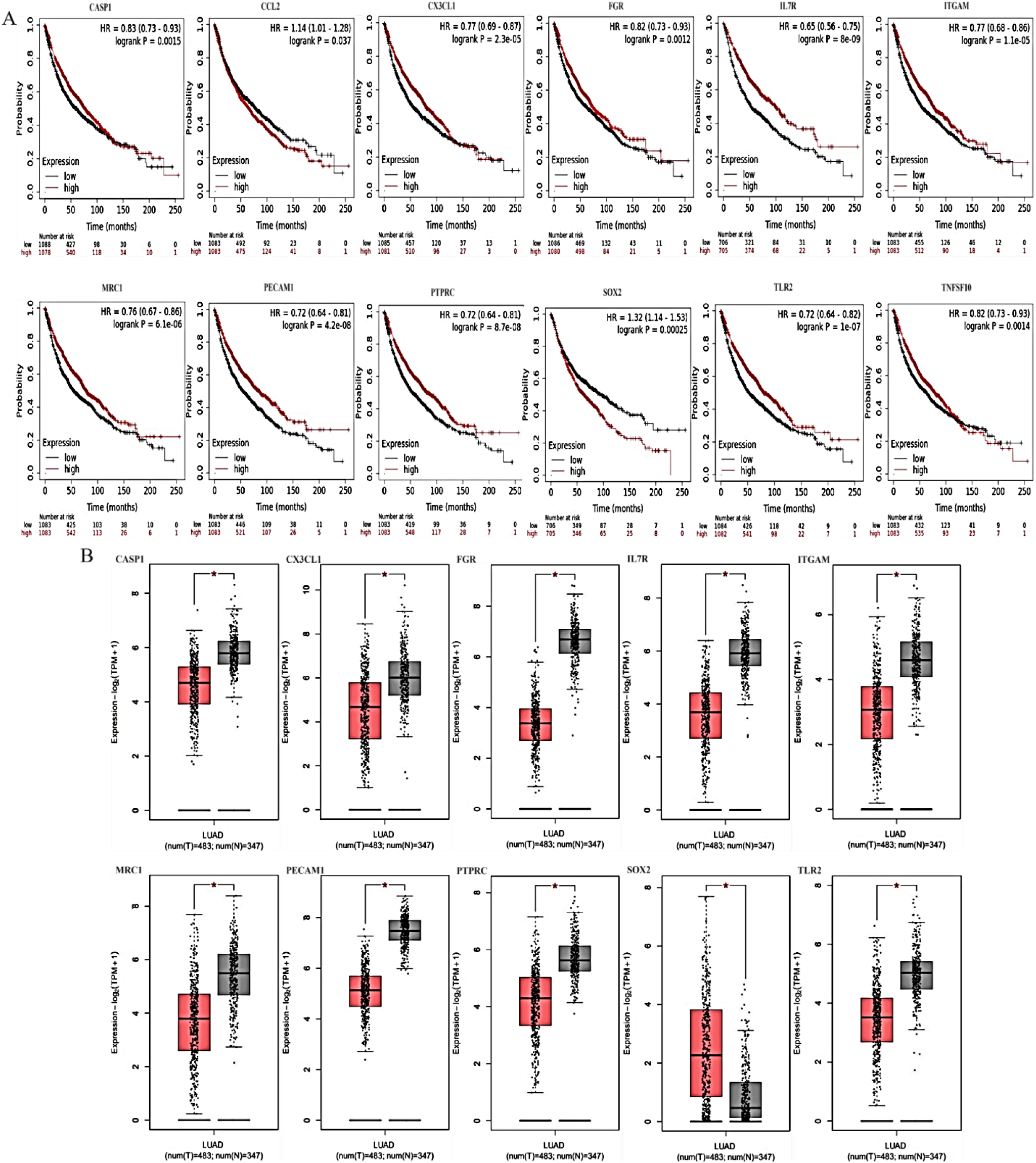
The association of mRNA expression with overall survival in patients with lung adenocarcinoma. (A) The Kaplan-Meier plots were generated by selecting the respective gene probe and the survival of all lung adenocarcinoma patients. The x-axis indicates the time of follow-up, and the y-axis indicates survival probability. Small vertical tick-marks indicate individual patients whose survival times have been right-censored. (B) mRNA expression of hub genes. mRNA expression of hub genes in LUAD tissue (red; n=483) and normal tissues (grey; n=347). CASP1, CX3CL1, FGR, IL7R, ITGAM, MRC1, PECAM1, PTPRC, SOX2, TLR2; * represents p value <0.05.

**Fig. 5.**
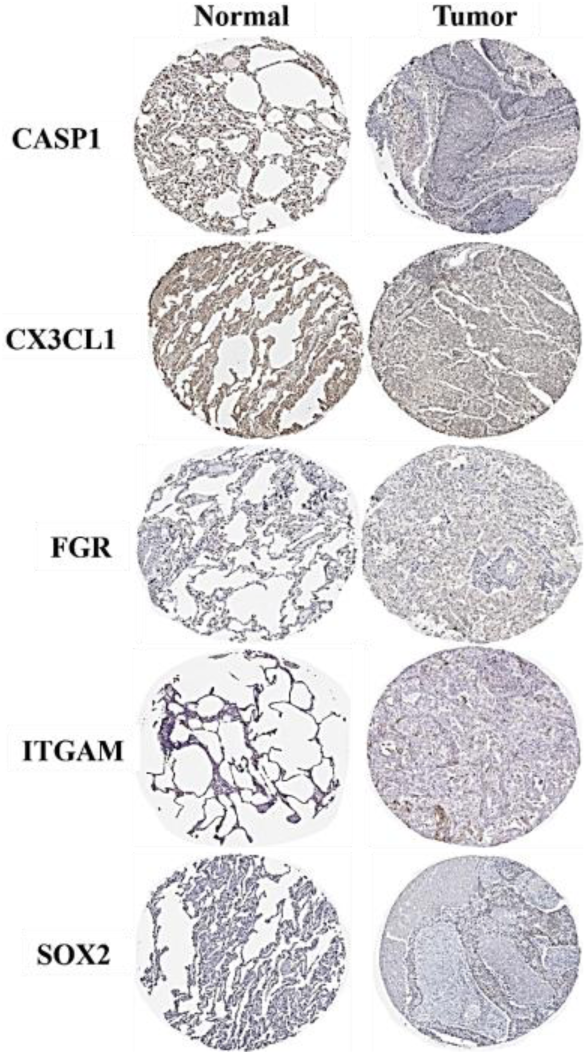
Validation of the hub genes from the HPA database. The representative image of suggested that the low expression of CASP1, CX3CL1, FGR, and ITGAM in lung adenocarcinoma tissue samples. However, SOX2 expression is higher in lung adenocarcinoma tissue samples in compare to normal lung tissue samples.

**Table 1:** Top 10 hub gene and their functions.

| Sl. No. | Gene Symbol | Official Full Name (NCBI) | Function |
| --- | --- | --- | --- |
| 1 | CASP1 | Caspase 1 | It plays a crucial role in cellular immunity as an initiator of the inflammatory response. Upon activation through the formation of an inflammasome complex, it triggers a pro-inflammatory reaction by cleaving the inflammatory cytokines IL1B and IL18, thereby releasing their mature forms, which participate in various inflammatory processes. |
| 2 | CX3CL1 | C-X3-C motif Chemokine Ligand 1 | This chemokine serves as a ligand for both CX3CR1 and the integrins ITGAV and ITGA4. The CX3CR1-CX3CL1 signaling pathway plays distinct roles in various tissue compartments, including immune response, inflammation, cell adhesion, and chemotaxis. |
| 3 | FGR | FGR Proto-Oncogene | This non-receptor tyrosine-protein kinase transmits signals from cell surface receptors lacking kinase activity and plays a role in regulating immune responses, including the functions of neutrophils, monocytes, macrophages, and mast cells. It also contributes to cytoskeleton remodeling in response to extracellular stimuli, as well as phagocytosis, cell adhesion, and migration. |
| 4 | IL7R | Interleukin 7 Receptor | It is a receptor for interleukin 7 (IL7). IL7R consists of an alpha/gamma chain heterodimer, with the gamma(c) chain primarily activating signal transduction and the alpha chain guiding specific signaling events. |
| 5 | ITGAM | Integrin Subunit Alpha M | Integrin ITGAM/ITGB2 is involved in various adhesive interactions of monocytes, macrophages, and granulocytes, and plays a role in mediating the uptake of complement-coated particles and pathogens. |
| 6 | MRC1 | Mannose Receptor C-type 1 | MRC1 facilitates the endocytosis of glycoproteins by macrophages and can bind both sulfated and non-sulfated polysaccharide chains. |
| 7 | PECAM1 | Platelet and Endothelial Cell Adhesion Molecule 1 | It is a cell adhesion molecule that is essential for leukocyte transendothelial migration (TEM) during most inflammatory conditions. |
| 8 | PTPRC | Protein Tyrosine Phosphatase Receptor Type C | It required for T-cell activation through the antigen receptor and acts as a positive regulator of T-cell coactivation upon binding to Dipeptidyl peptidase-4 (DPP4). |
| 9 | SOX2 | SRY-box transcription factor 2 | It plays a role in regulating embryonic development and determining cell fate. The gene product is essential for stem cell maintenance in the central nervous system. SOX2 forms a trimeric complex with OCT4 on DNA, regulating the expression of several genes involved in embryonic development, including YES1, FGF4, UTF1, and ZFP206. |
| 10 | TLR2 | Toll Like Receptor 2 | It works with LY96 to mediate the innate immune response to bacterial lipoproteins and other microbial cell wall components. Additionally, it collaborates with TLR1 or TLR6 to facilitate the innate immune response to bacterial lipoproteins or lipopeptides. |

### 3.6 Interactive network analysis of TFs

In order to get the functional network of SOX2, we used the TRRUST database and got the top 20 SOX2 targets. Here, we found that MMP3 and two members of the superfamily of ATP-binding cassette (ABC) transporters, ABCC3 and ABCC6, are activated by SOX2 [49,50]. Additionally, SOX2 activates DKK1, a protein that binds to the LRP6 co-receptor to block Wnt signaling dependent on beta-catenin. Many human cancers have been reported to have elevated DKK1 expression, and its protein may boost cancer cell line proliferation, invasion, and growth [51–53]. However, BMP4 is repressed by the action of SOX2. Studies suggested that SOX2 transcriptionally represses the BMP4 promoter, and hence, in LSCC, BMP4 functions as a tumor suppressor. However, SOX2 suppressing BMP4 transcription promotes cell proliferation [54].

### 3.7 SOX2-EZH2 exhibit a positive correlation in LUAD prognosis

Based on the TF-gene regulatory network analysis, SOX2 gene directly interacts with a pool of transcription factors, including EZH2, ZNF71, POU5F1B, HIC1, CBX8, CTBP2, BCL11B, and KLF9 (Figure 6A). In cancer, EZH2 is frequently overexpressed and repress tumor suppressor gene transcription and contributes in oncogenic signaling facilitating tumor development and cancer progression/metastasis. The regression analysis showed that, the relevant coefficients for SOX2 and EZH2 were high i.e., Pearson’s correlations are 0.73 and 0.66, respectively (Figure 6B). The authenticity of connection was verified by utilizing UCSC Xena to examine patients with lung cancer data from the TCGA database. These findings suggested a potential connection between SOX2 and EZH2-mediated gene regulation in lung cancer. The SOX2 gene was implicated as possibly co-expressing with EZH2 (Figure 6C). Using cBioPortal, data mining was done in a lung cancer cohort by querying the public TCGA lung adenocarcinoma samples to learn more about the underlying positive correlation of SOX2 mRNA and EZH2 mRNA samples (Figure 6D). The OncoDB database confirmed the positive connection between the transcript expression of SOX2 and EZH2 (Figure 6E).

**Fig. 6.**
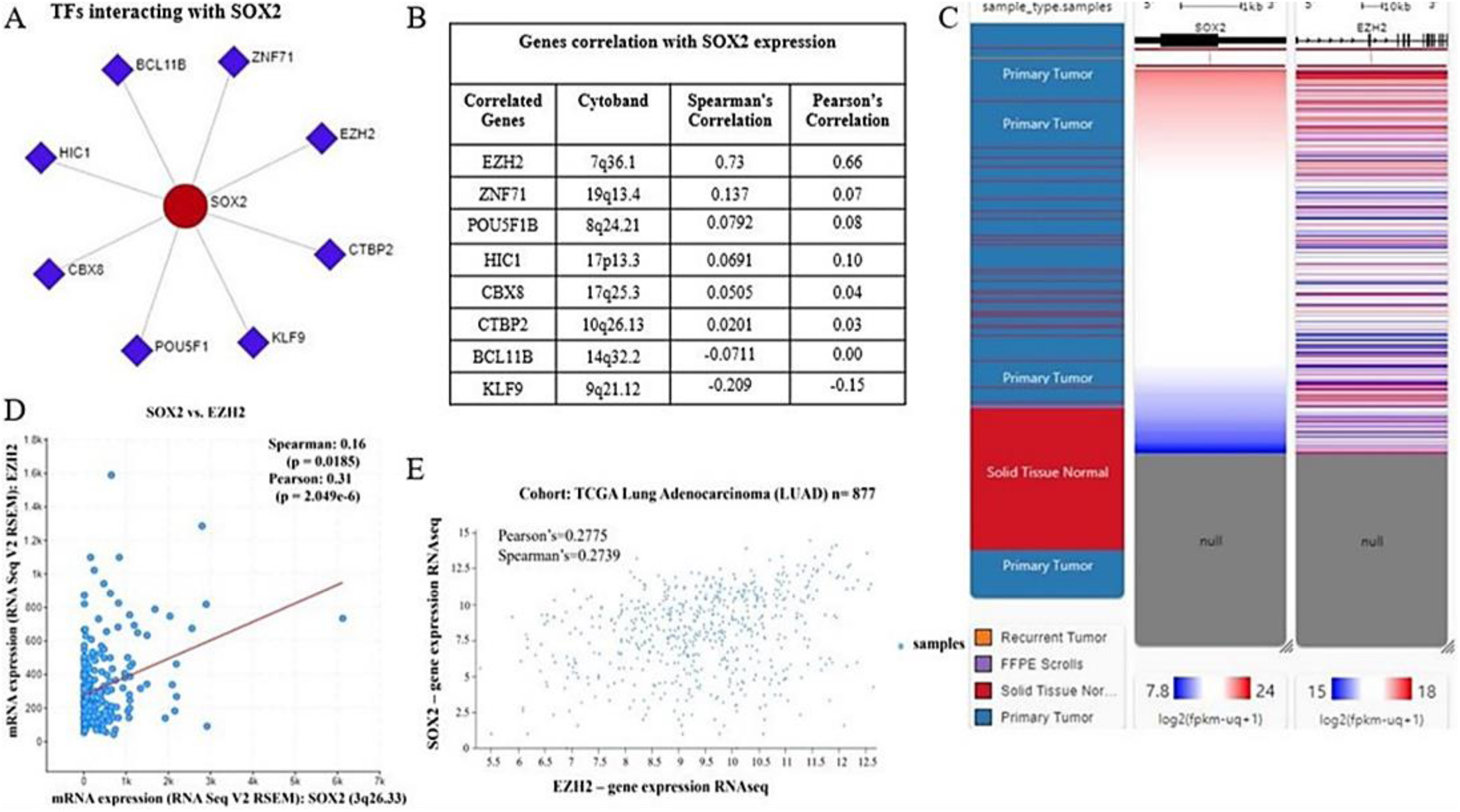
SOX2 gene expression is associated with other TFs in lung cancer. (A) Interaction of SOX2 gene regulatory networks incorporating transcription factors such as EZH2, CTBP2, KLF9, POU5F1, CBX8, HIC1, BCL11B, and ZNF71. (B) The SOX2 gene positively associated with SOX2 transcript level based on TCGA among ∼230 patients with lung cancer. (C) Through regression analysis, it was determined that SOX2 and EZH2 were highly correlated. (D) Data mining in the OncoDB database further confirmed the positive correlation of SOX2 and EZH2 mRNA expression. (E) A heatmap derived from University of California, Santa Cruz (UCSC) Xena revealed the SOX2 and EZH2 mRNA expression levels among lung cancer subtypes in TCGA database. TCGA, The Cancer Genome Atlas; SOX2, Sex determining region Y-box 2; EZH2, Enhancer of zeste homolog 2.

### 3.8 SOX2 expression is relatively very high in lung cancer cells, indicating its involvement in cancer progression

Following our understanding of DEGs expression by applying bioinformatics, we conducted qRT-PCR analysis. We compared the corresponding mRNA expression in A549 and NCI-H522 cells to that of HaCaT, and β-actin was taken as an internal control. Our findings aligned with the previously mentioned findings. Here, we found that SOX2 expression in A549 and NCI-H522 cells is significantly higher than HaCaT cells while expression of CASP1, CCL2, CX3CL1, FGR, IL7R, ITGAM, MRC1, PECAM1, PTPRC, TLR2 is relatively low (Figure 7). Here also, we have observed that SOX2 is the only gene that is showing higher expression among all the ten genes. Hence, we proceed to protein expression of SOX2. Western blot analysis suggested that, A549 and NCI-H522 cells had significantly high expression of SOX2, which aligns with the result of increased mRNA expression and supports the findings of the qRT-PCR data (Figure 8).

**Fig. 7.**
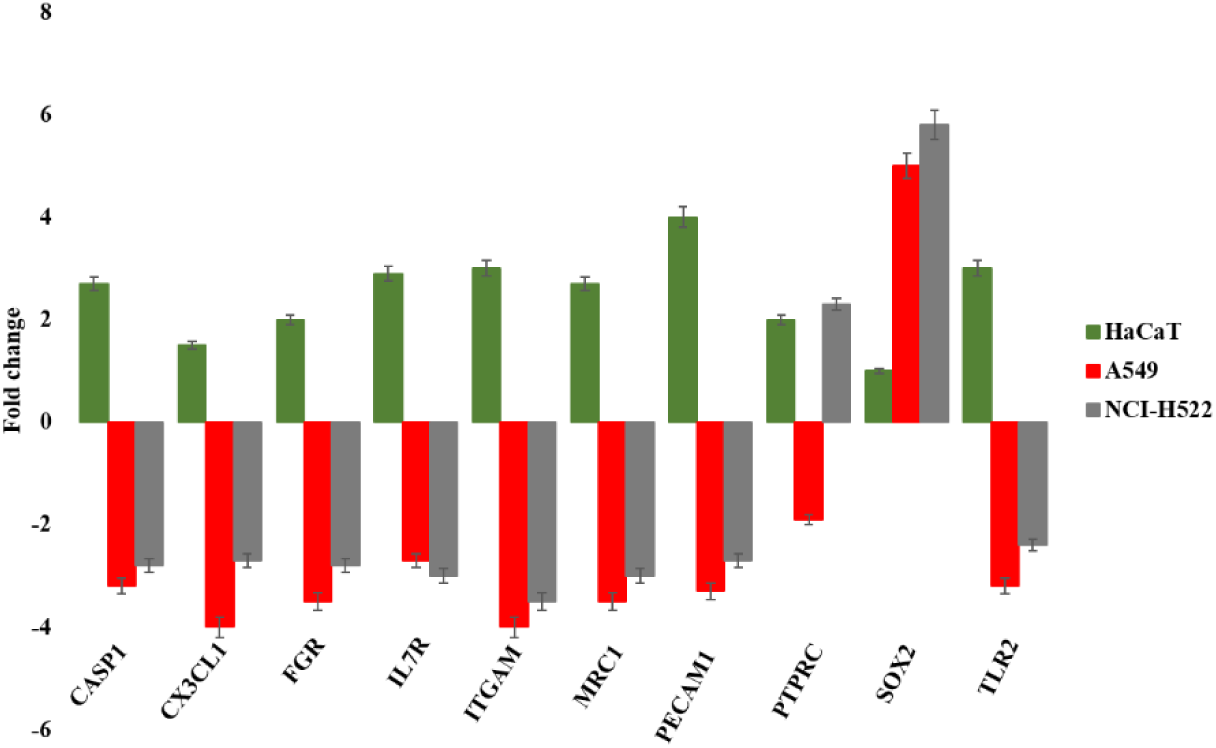
Expression profiling of CASP1, CX3CL1, FGR, IL7R, ITGAM, MRC1, PECAM1, PTPRC, SOX2, and TLR2 in A549 and NCI-H522 cells. Relative mRNA expression level of SOX2 in comparison to HaCaT cells (control).

**Fig. 8.**
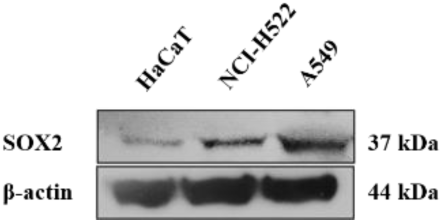
Western blot analyses of SOX2 protein in HaCaT, A549, and NCI-H522 cells. It is apparent that in normal skin cell line (HaCaT) there is no expression of SOX2.

### 3.9 The repurposed drugs interact with EZH2

FTSite server identified three functional sites (Figure 9A & B). Molecular docking using AutoDock 4.2.6 was performed to study the molecular interactions between EZH2 and repurposed drugs listed on Table 3. Based on the docking study, the drug GSK343 exhibited the highest binding affinity for EZH2, demonstrating superior binding energy compared to other drugs (Figure 10A & B). GLY A: 623, TRP A: 624, ALA A: 622, ASN A: 688, HIS A: 689, ALA A: 687, PHE A: 665, ARG A: 685, TRP A: 113, SER A: 112, THR A: 678, VAL A: 657, TYR A: 658, TYR A: 111, TYR A: 661, and ILE A:109 were crucial in bond formation between EZH2 and GSK343 (Figure 10C).

**Fig. 9.**
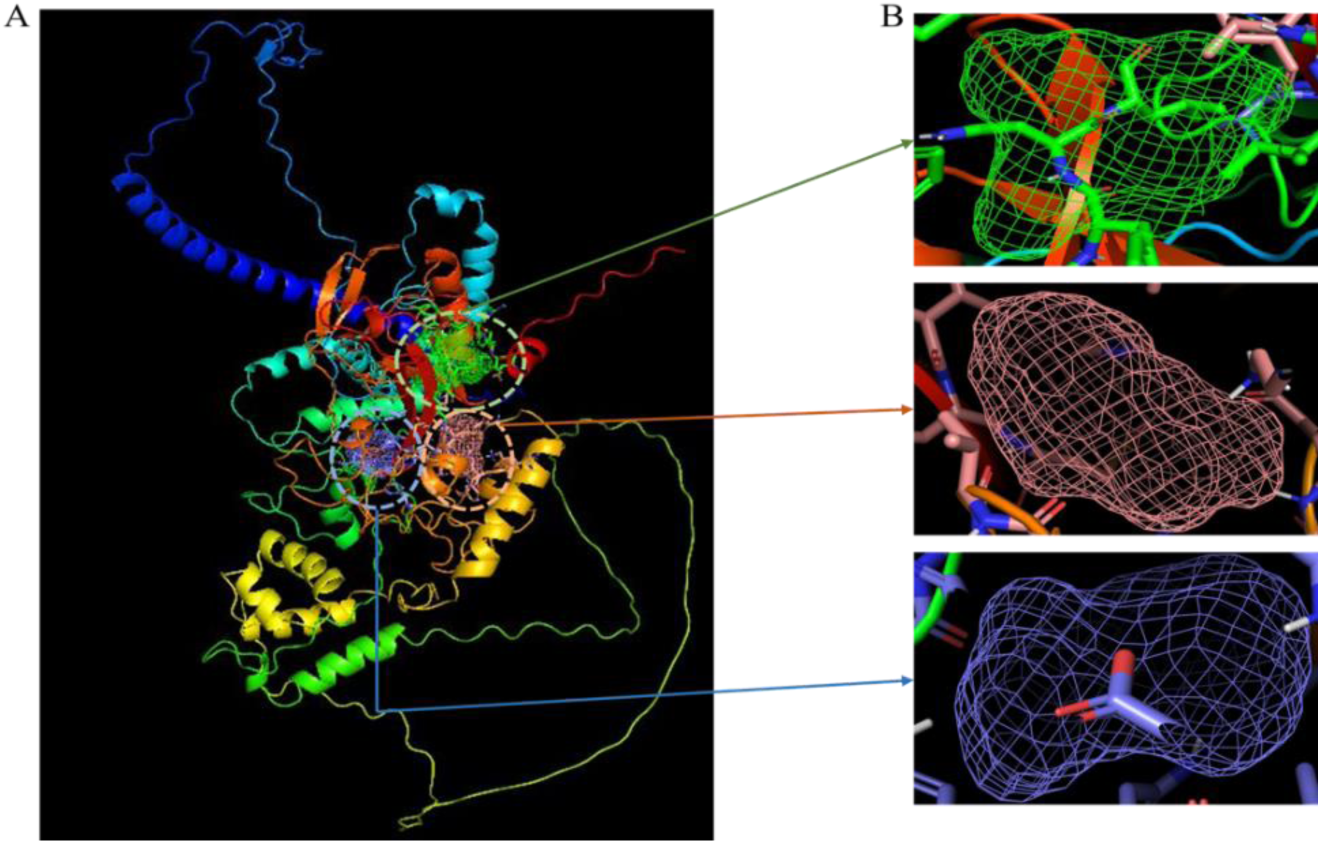
Three-dimensional structure (3D) of EZH2. (A) Three different functional sites were observed using FTSite (https://ftsite.bu.edu/) and visualized in PyMol, taking PDB id: 5HYN as input. The dotted circles represent the functional site of the respective proteins. (B) Zoomed and centered view of three different active sites (Green, Orange, and Blue).

**Fig. 10.**
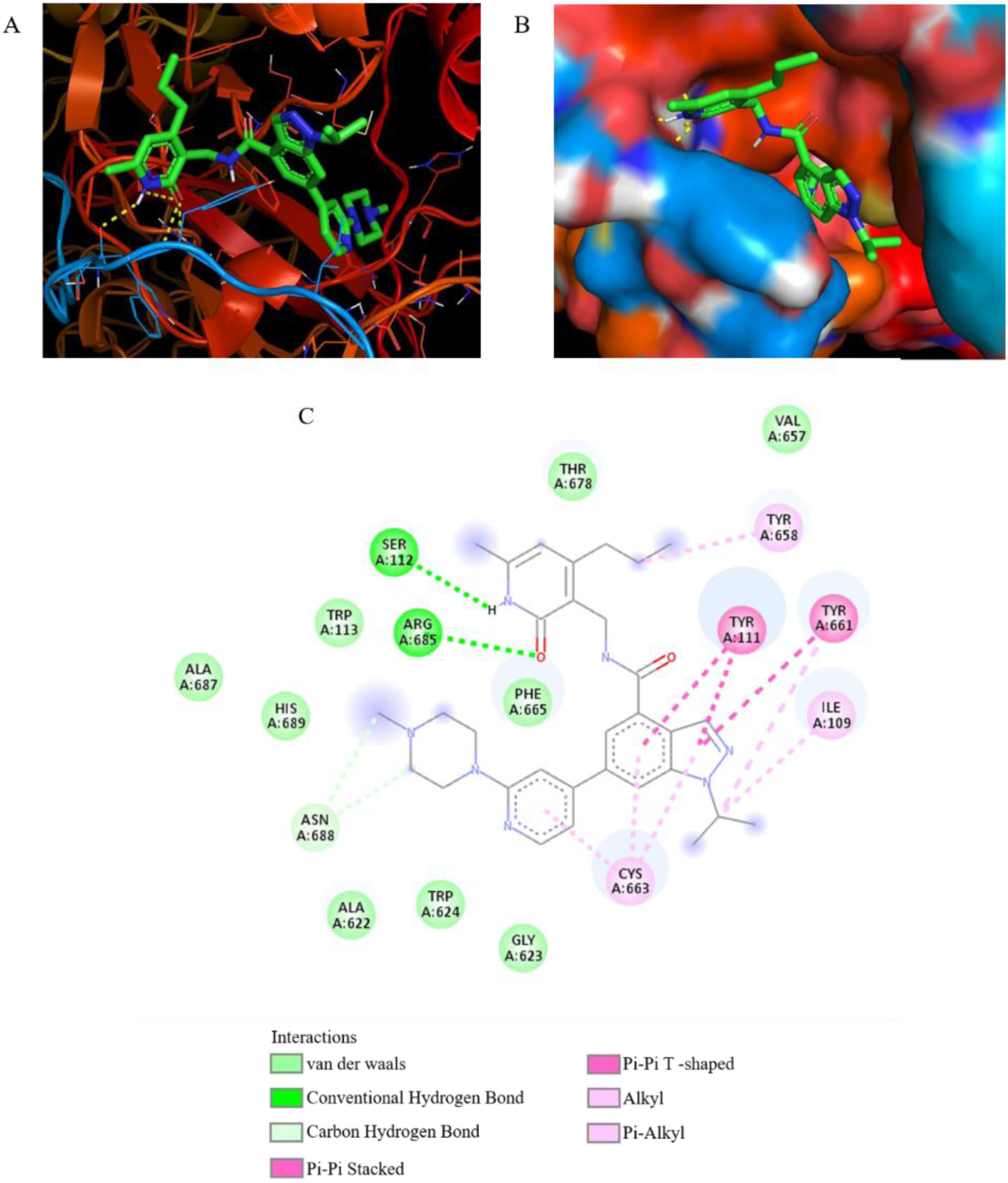
Molecular docking of EZH2 with inhibitors. (A) GSK343 binds at the active site of EZH2. (B) Zoomed view of GSK343 in the pocket of EZH2. (C) For visualization, BIOVIA Discovery Studio Visualizer used and 2D structure was generated and showcase the bonds between EZH2 and GSK343 that are GLY A: 623, TRP A: 624, ALA A: 622, ASN A: 688, HIS A: 689, ALA A: 687, PHE A: 665, ARG A: 685, TRP A: 113, SER A: 112, THR A: 678, VAL A: 657, TYR A: 658, TYR A: 111, TYR A: 661, and ILE A:109.

**Table 2:**
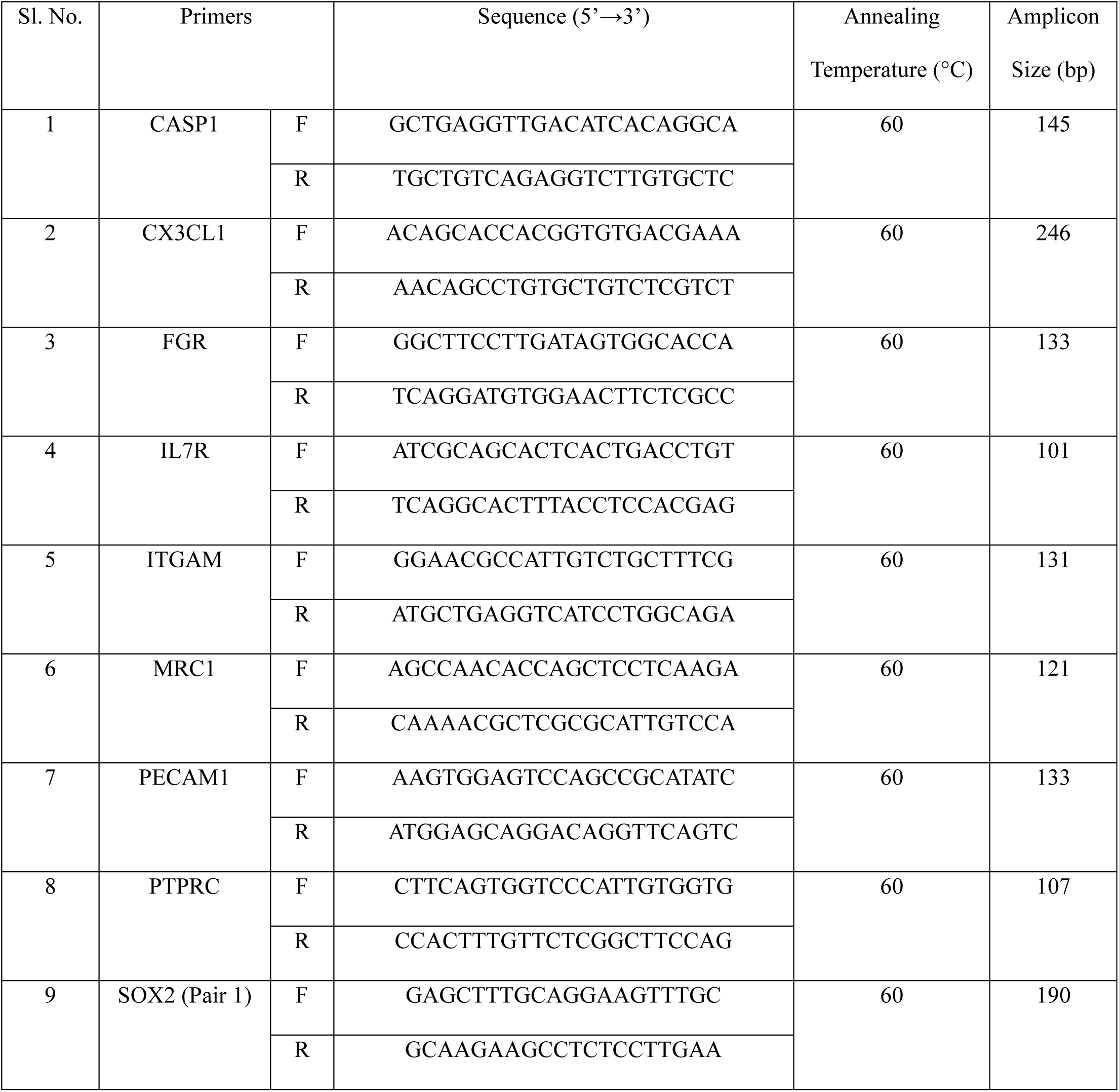
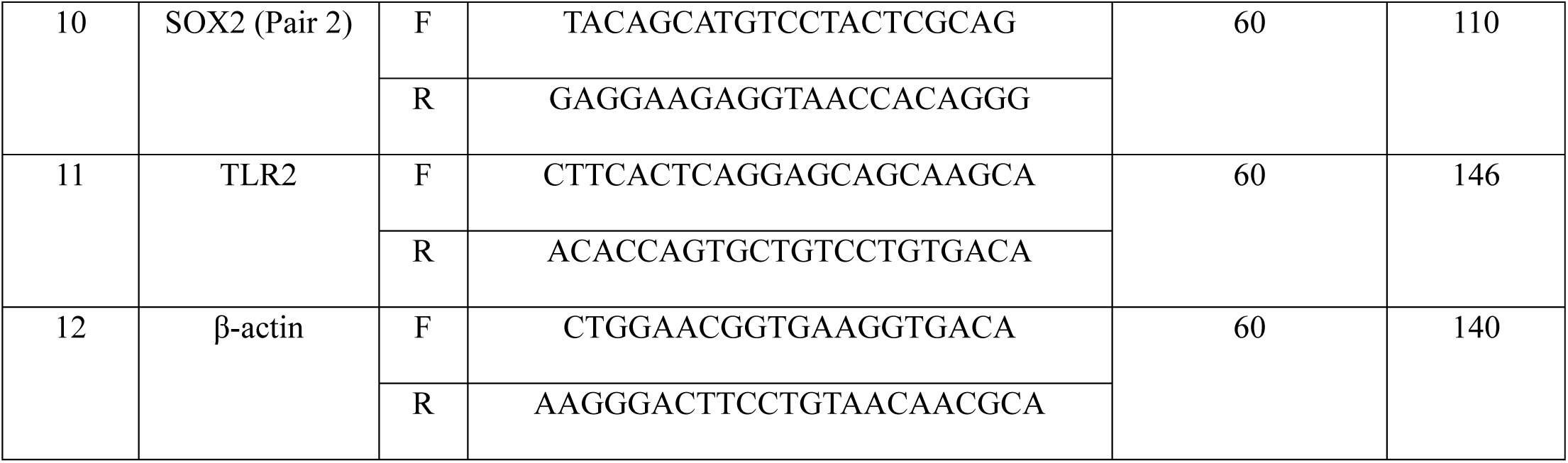
List of primers used for this study. F= Forward, R=Reverse.

**Table 3:**
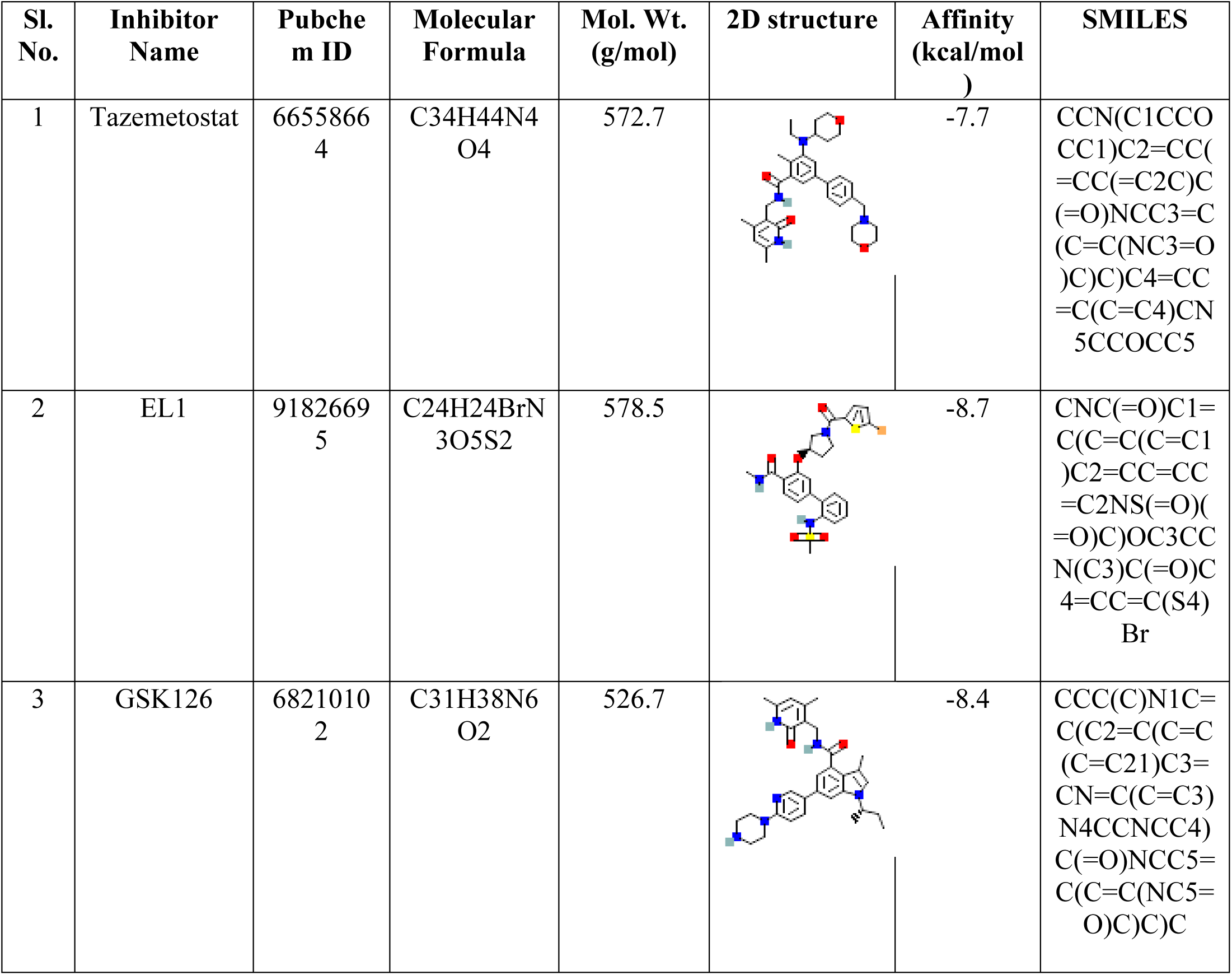
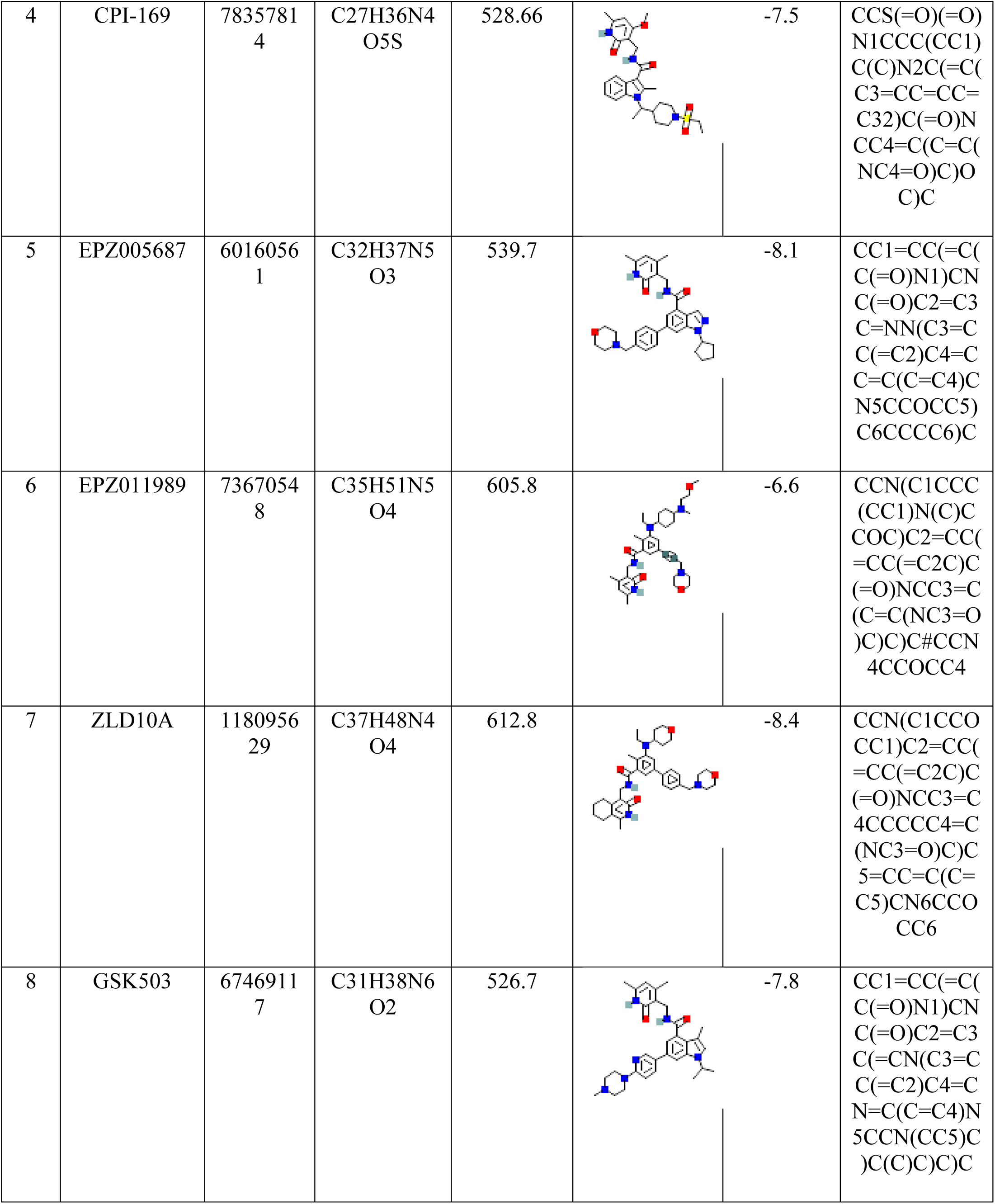
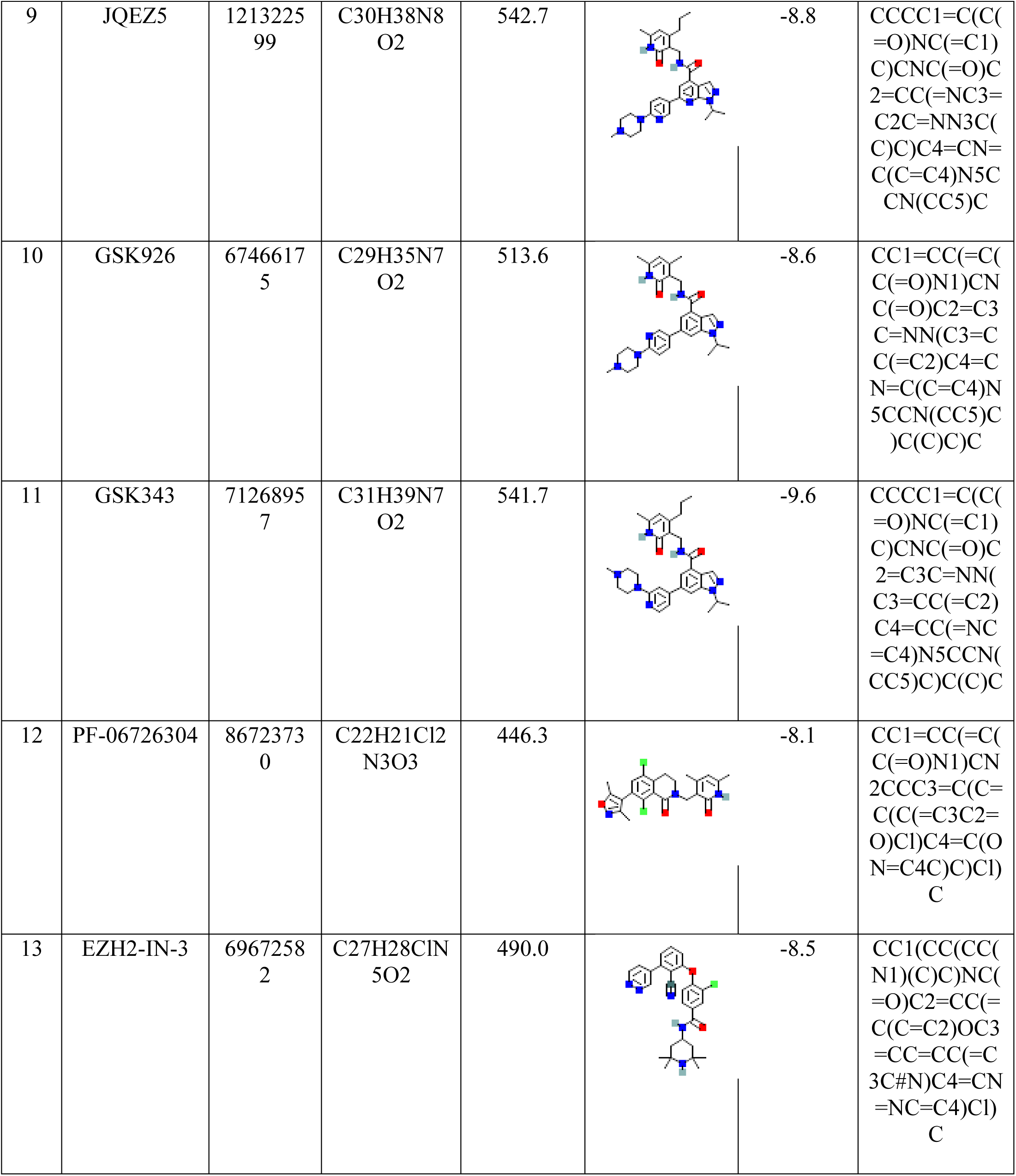
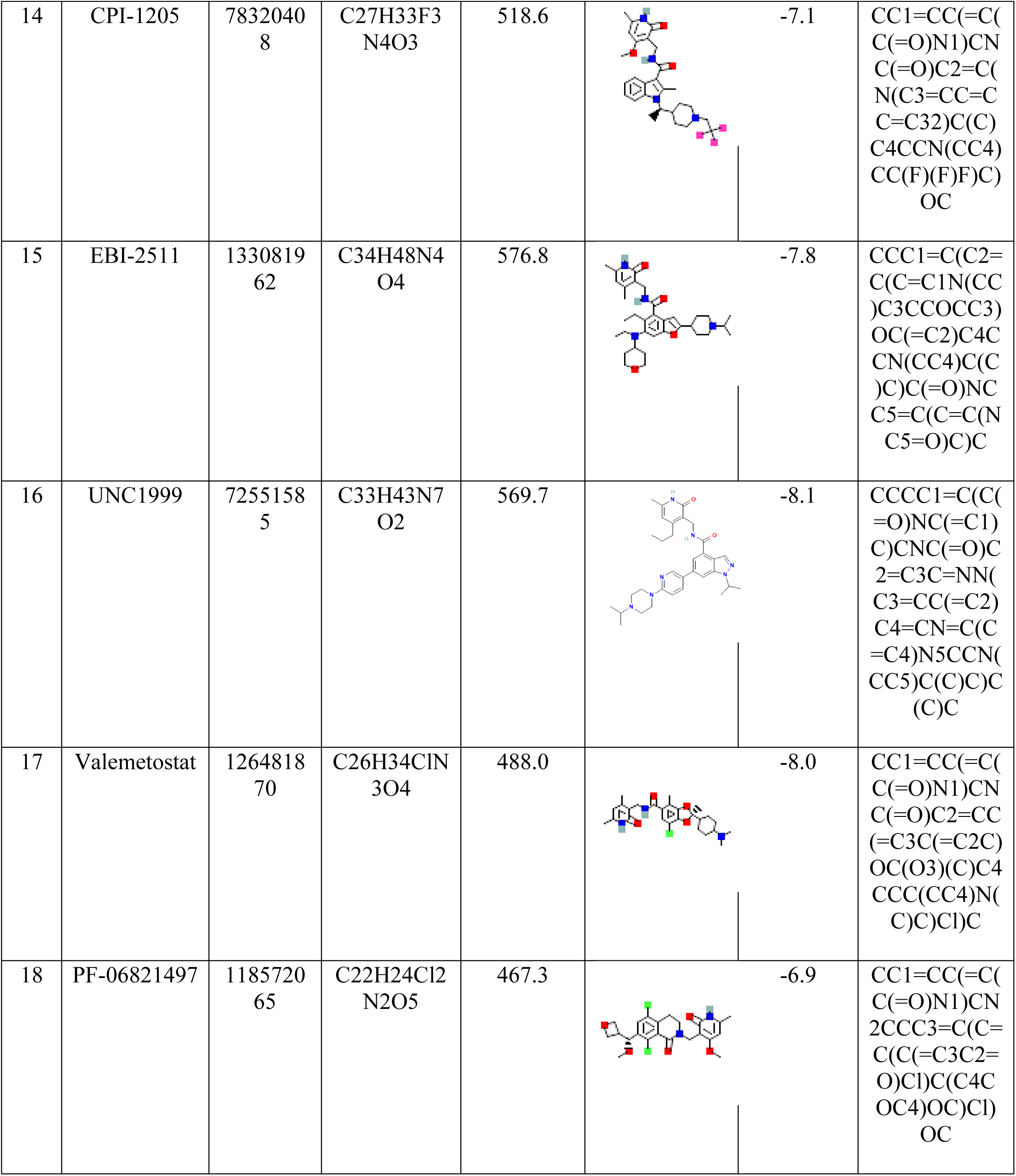
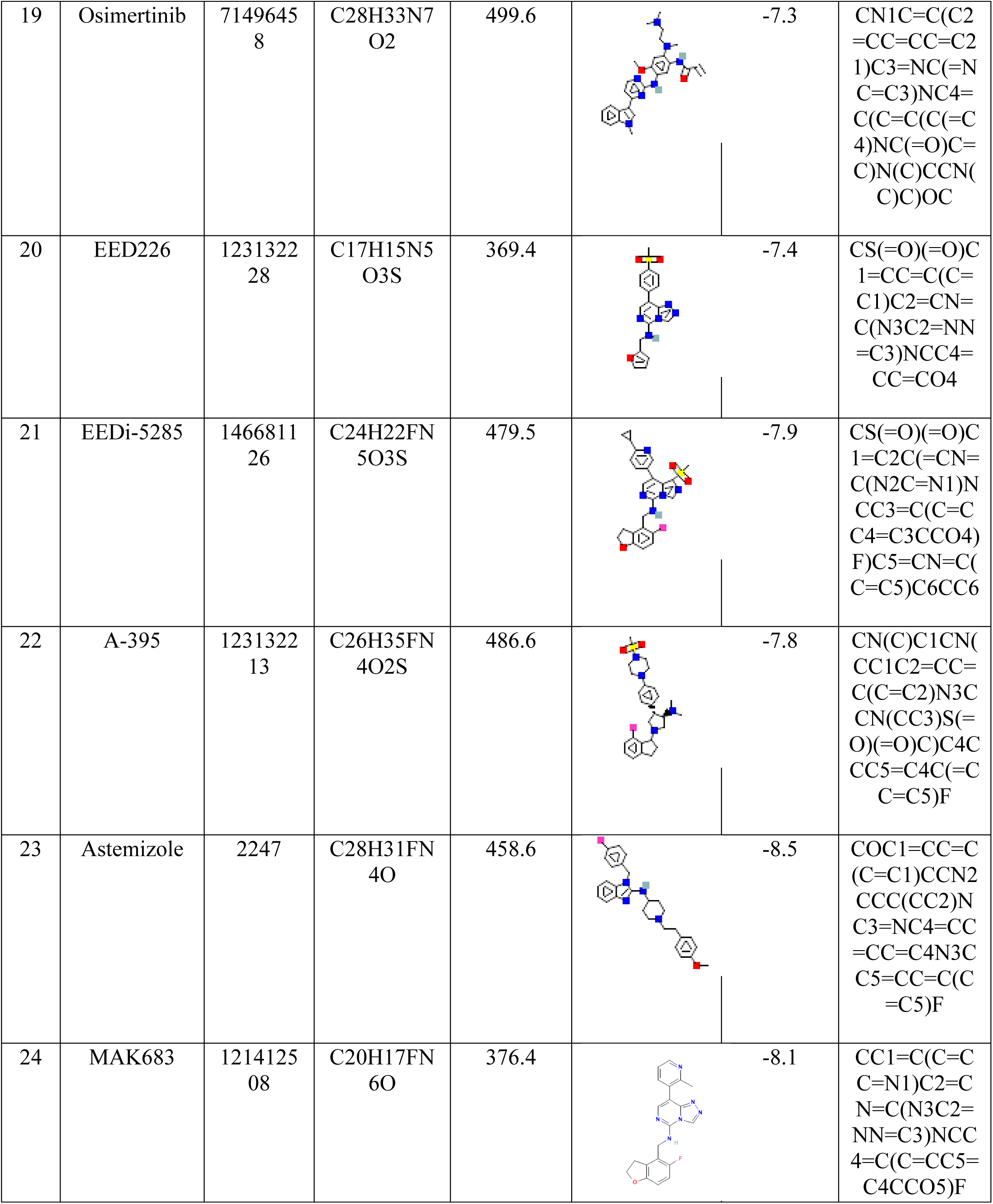
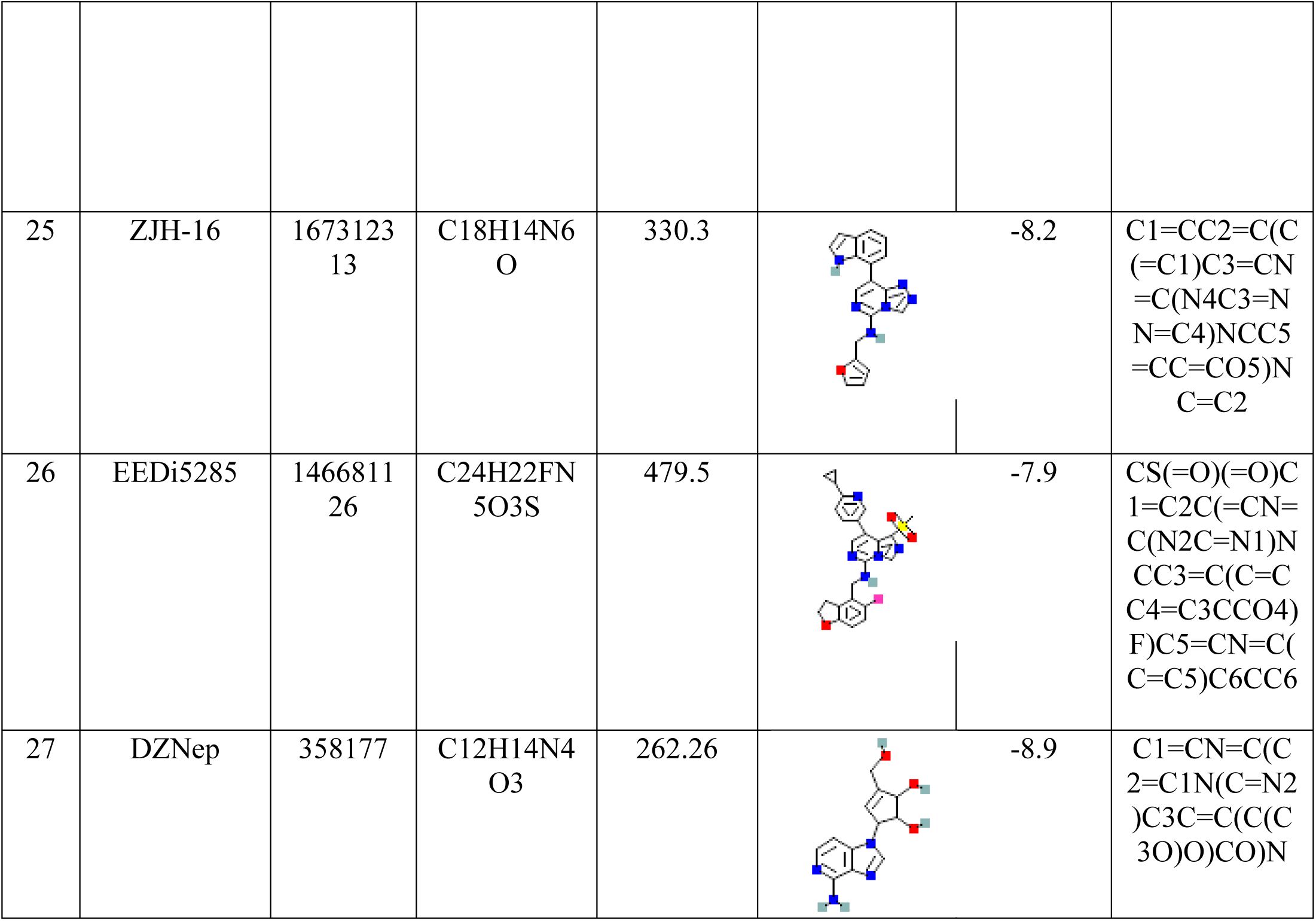
The binding energy of inhibitors bound to EZH2.

## 4 Discussion

Efforts to manage lung cancer is not satisfactory, the prognosis is not justifiable. Late detection, due to a lack of reliable early diagnosis, has led to high mortality rate. While recent years have witnessed the emergence of targeted gene therapy as a novel treatment approach, supported by some successful experiments, it is yet unknown what is/are molecular mechanism(s) underlying pathophysiology of lung cancer. Consequently, it is critical to investigate the key molecule(s) linked molecular pathways accountable initiation and advancement of lung cancer and to find useful early tumor indicators [55–57].

Advanced technologies, including bioinformatics analysis and the examination of microarray and sequencing data, play a crucial role in uncovering key genetic alterations in cancer. In this work, we screened DEGs in two GEO datasets (GSE19188, GSE68465) using several bioinformatics methods, and we found 795 common DEGs in total. Among these, 337 genes were commonly upregulated, while 458 genes were commonly downregulated. Subsequent gene ontology and KEGG pathway enrichment analyses indicated significant enrichment of DEGs in cancer-related pathways. Further analysis identified 12 candidate genes with significance in survival analysis. Notably, lower expression levels of CASP1, CCL2, CX3CL1, FGR, IL7R, ITGAM, MRC1, PECAM1, PTPRC, TLR2, TNFSF10 were significantly associated with poor OS in lung cancer patients.

In contrast, higher expression levels of SOX2 were also significantly linked to poor OS in lung cancer patients. Validation of these findings using GEPIA2 and HPA databases revealed that in LUAD samples, SOX2 mRNA expression was notably higher than in normal samples. Conversely, CASP1, CX3CL1, FGR, IL7R, ITGAM, MRC1, PECAM1, PTPRC, and TLR2 genes exhibited higher expression levels in normal samples compared to LUAD samples. Protein expression data from the HPA database supported the overexpression of SOX2 in LUAD samples, correlating with lung cancer prognosis. To further validate the observed SOX2 expression trends at both the mRNA and protein levels, qRT-PCR and western blot analyses were conducted in A549 and NCI-H522 cells. The findings consistently showed that lung cancer cells expressed SOX2 more frequently than normal cells in both mRNA and protein levels.

SOX2 and EZH2 are proteins with pivotal roles in diverse cellular processes, regulating gene expression/repression during embryonic development. Anomalies in the expression of and regulation by these proteins have been linked to cancer. SOX2 is a transcription factor essential for preserving the pluripotency of embryonic stem cells. It also plays a complex role in controlling developmental pathways. The overexpression of SOX2 has been reported in several cancer types, including esophageal, breast, ovarian, and lung cancers. In certain instances, elevated SOX2 levels are associated with cancer stem cell properties, contributing to tumor initiation, progression, and resistance to therapeutic interventions [16,17,58].

On the other side, EZH2 functions as a catalytic subunit of the PRC2 and it is a histone methyltransferase. It is involved in epigenetic regulation of gene expression by adding methyl groups at lysine 27 of histone H3 (H3K27), resulting in chromatin inactivation and gene silencing. Many cancers, such as prostate, breast, lung, and hematological malignancies, commonly overexpress EZH2. This overexpression often correlates with more aggressive tumor behavior, poor prognosis, and resistance to conventional therapies. The epigenetic alterations mediated by EZH2 contribute to silencing tumor-suppressor genes [59].

The interaction between SOX2 and EZH2 in the context of cancer progression is an evolving area of research, with emerging evidence suggesting potential collaborative roles for these two proteins in certain cancers. In specific cancer types, co-expression of SOX2 and EZH2 is observed, indicating elevated levels of both proteins within the same tumor cells. Despite differing mechanisms, both SOX2 and EZH2 impact gene expression. SOX2, functioning as a transcription factor, directly regulates the expression of specific genes. The coordinated activities of SOX2 and EZH2 may contribute to the acquisition of aggressive phenotypes during cancer progression. In case of prostate cancer EZH2 and SOX2 has been shown to correlated with tumor progression and high Gleason score [60–62]. The coordination of EZH2 and SOX2 plays important role in human neural fate decision [63].

The molecular interaction between EZH2 (PDB ID: 5HYN) with their repurposed drugs was studied by performing molecular docking using AutoDock. The docking study revealed that the drug GSK343 showed the highest binding affinity for EZH2, compared to the other listed drugs. Table 3 represents the binding energy of target proteins in the active region with their corresponding drugs. As shown above, GSK343 binds to EZH2 with higher binding energy (Figure 10), indicating this drug can be more effective in reversing the expression of EZH2 along with SOX2 and can be implicated in lung adenocarcinoma treatment. Thus, it is hypothesized that these GSK343 could nullify the effect of EZH2-mediated regulations over SOX2 promoting lung tumorigenesis.

Utilizing the similar methodology and employing similar bioinformatic tools we explored DEGs profiles in Ovarian Cancer (OvCa) tissue samples and identified six key hub genes, including BUB1B, CCNA2, MAD2L1, PRC1, TRIP13 and ZWINT. DEGs analysis of various molecular subtypes implicated the significance of the hub genes in aggressive subtypes of OvCa [35].

## 5 Conclusion

This work revealed the potential involvement of the SOX2-EZH2 axis in the initiation and metastasis of lung cancer. Beyond its role as a pluripotent factor, SOX2 and chromatin repressor partner EZH2 would be novel biomarkers and therapeutic targets for individuals with lung cancer. Furthermore, an inhibitor, GSK343 exhibited a promising therapeutic compound that can imply to inhibit EZH2. Overall, this study analyzed and presented a precise methodology for identifying and prioritizing the most effective candidate genes associated with lung cancer prognosis, and could be a model for identifying new biomarker(s) for other cancers.

## Key Points

- Lung adenocarcinoma (LUAD) accounts for approximately 40-50% of all lung cancer cases; hence, for better treatment of lung cancer deep research functional analysis of genes is essential.
- Using database for LUAD patient sample containing information for both normal and disease states, we identified that, genes SOX2 and EZH2 are involved in development and progression of LUAD.
- EZH2 trimethylates H3K27 (forming H3K27me3) and repress genes. EZH2 can be targeted by the drug, GSK343, for eventually hampering of the SOX2-EZH2 cooperation, for better treatment of LUAD.

## Data Availability Statement

All data generated or analyzed during this study are included in this article. All data available for the audience and further inquiries can be directed to the corresponding author.

## Author Contributions

SKP and RS conceptualized the project. Niharika and A. Roy performed all the data search and experiments. Niharika designed and analyzed the experiments and wrote the manuscript. RS checked the manuscript and SKP edited the final manuscript.

## Ethics approval and consent to participate

Not applicable.

**Niharika** is a PhD student at Epigenetics & Cancer Biology Lab, Department of Life Science at National Institute of Technology Rourkela. Her research focuses on epigenetic mechanisms that regulate pluripotency-inducing transcription factors in cancer *vis-a-vis*, utilizing molecular biology and genomics approaches.

**Ankan Roy** is a PhD student at Epigenetics & Cancer Biology Lab, Department of Life Science at National Institute of Technology Rourkela. His research interests inclue lipidraft mediated destabilization of epiegenetic choreography in cancer.

**Ratan Sadhukhan is a staff scientist, who had his PhD in biophysics** with huge experience in molecular biology, bioinformatics and cancer. Current research interest is on DNA damage repair, cell biology and oncology.

**Samir Kumar Patra** is a Professor and former Head of the Department of Life Science at the National Institute of Technology Rourkela. A biochemist and molecular cell biologist with well over thirty years of research experience on biochemistry, molecular biology, epigenetics and cancer; broad area of research includes chromatin modifications, cell signaling, mechanotransductiuon, liquid-liquid phase separation, gene expression analyses and oncology.

